# Uncertainty quantification of reference based cellular deconvolution algorithms

**DOI:** 10.1101/2022.06.15.496235

**Authors:** Dorothea Seiler Vellame, Gemma Shireby, Ailsa MacCalman, Emma L Dempster, Joe Burrage, Tyler Gorrie-Stone, Leonard S Schalkwyk, Jonathan Mill, Eilis Hannon

## Abstract

The majority of epigenetic epidemiology studies to date have generated genome-wide profiles from bulk tissues (e.g. whole blood) however these are vulnerable to confounding from variation in cellular composition. Proxies for cellular composition can be mathematically derived from the bulk tissue profiles using a deconvolution algorithm however, there is no method to assess the validity of these estimates for a dataset where the true cellular proportions are unknown. In this study, we describe, validate and characterise a sample level accuracy metric for derived cellular heterogeneity variables. The CETYGO score captures the deviation between a sample’s DNAm profile and its expected profile given the estimated cellular proportions and cell type reference profiles.We demonstrate that the CETYGO score consistently distinguishes inaccurate and incomplete deconvolutions when applied to reconstructed whole blood profiles. By applying our novel metric to > 6,300 empirical whole blood profiles, we find that estimating accurate cellular composition is influenced by both technical and biological variation. In particular, we show that when using the standard reference panel for whole blood, less accurate estimates are generated for females, neonates, older individuals and smokers. Our results highlight the utility of a metric to assess the accuracy of cellular deconvolution, and describe how it can enhance studies of DNA methylation that are reliant on statistical proxies for cellular heterogeneity. To facilitate incorporating our methodology into existing pipelines, we have made it freely available as an R package (https://github.com/ds420/CETYGO).

## Introduction

Due to the dynamic nature of the epigenome and its plasticity in response to environmental exposures (Hannon et al., 2018, Joehanes et al., 2016, Tobi et al., 2014, Gruzieva et al., 2017), there is increasing interest in the role it plays in the aetiology of disease (Murphy and Mill, 2014). However, this very facet of the epigenome makes epigenetic epidemiology studies inherently more complex to design and liable to confounding compared to studies of DNA sequence variation (Heijmans and Mill, 2012, Relton and Davey Smith, 2010). One major difference is that an individual’s genetic sequence is identical in all cells, and therefore it does not matter from which tissue DNA is isolated prior to genotyping. In contrast, the epigenome orchestrates gene expression changes that underpin cellular differentiation, consequently, cell types can be defined by their epigenetic profiles (Stunnenberg et al., 2016, Roadmap Epigenomics Consortium et al., 2015). It has previously been shown that variation between cell types is greater than inter-individual variation within a cell type (Hannon et al., 2021b, Shanthikumar et al., 2021).

The majority of studies to date have focused on a single epigenetic modification, DNA methylation, and generated genome-wide profiles from bulk tissues (e.g. whole blood) using high throughput microarrays (Campagna et al., 2021). A critical challenge in these studies is that bulk tissue is a heterogeneous mix of different cell types. The epigenetic profile of a bulk tissue is the average across the profiles of the constituent cell types. If the composition of these cell types, specifically the proportions of each cell type, varies across the population under study, and varies in a manner that correlates with the outcome of interest, this will lead to false positive associations at sites in the genome that differ between cell types (Jaffe and Irizarry, 2014, Liu et al., 2013). As a result, epigenome-wide association analyses routinely include quantitative covariates that capture the heterogeneity in cellular composition across a dataset. As experimentally derived cell counts are often unavailable, proxies for cellular composition can be derived from the bulk tissue profile using a deconvolution algorithm. The goal of these statistical methodologies is to generate a series of continuous variables that reflect the underlying cellular heterogeneity of each sample. Deconvolution algorithms can be separated into two classes. Firstly, supervised methods that incorporate reference profiles for relevant cell types - generated from purified cell populations - and estimate proportions for this specified set of cell types (known as reference-based algorithms)(Houseman et al., 2012, Newman et al., 2015, Accomando et al., 2014, Guintivano et al., 2013, Teschendorff et al., 2017). Secondly, those that do not use any reference data and generate an unlimited set of variables that are not directly attributed to any particular cell type (known as reference-free algorithms)(Houseman et al., 2014, Leek and Storey, 2007, Rahmani et al., 2019, Zou et al., 2014).

In tissues for which reference profiles are available, reference based deconvolution algorithms are most commonly used, likely due to the ease of interpretation. Specifically the constrained projection methodology proposed by Houseman, often referred to as “Houseman’s method”, is normally used. There have been a number of studies that have aimed to validate the application of these methods by testing their performance against experimentally or computationally derived “bulk” profiles of fixed cellular compositions (Koestler et al., 2013, Salas et al., 2018). These have primarily focused on the prediction of the major blood cell types from whole blood. Typically, accuracy is reported at the group level, i.e. a single correlation or error statistic across a number of samples, which is then assumed to be representative for all future applications. In prediction modelling, great attention is paid to ensuring that the training data is representative of the testing data to so that the predictions are valid. The vast majority of whole blood epigenetic studies use the same reference dataset generated from six adult males to determine cellular composition, regardless of the age, sex, ethnicity, or disease status characteristics, with little consideration given to whether it is representative of the cohort being tested. Mathematically, there is nothing to prevent a deconvolution algorithm, based on any reference panel of cell types, from being applied to a profile generated from any bulk tissue. As an extreme example, we could input data derived from brain tissue to a model that outputs estimates of the composition of blood cell types and obtain values, due to the mathematical constraints, that are plausible (i.e. between 0 and 1). In a less extreme example, it is unknown how important demographic features (e.g. age, sex, or ethnicity) of the samples in the reference panel affect prediction in samples characterised by different demographics. Currently, there is no method to assess the validity of cellular composition estimates for a single sample, or indeed, a dataset where the true cellular proportions are unknown. If the quality of the deconvolution varies either, across studies or within a study, then the utility of these variables as confounders needs to be reconsidered. This could be especially problematic if the accuracy of the deconvolution is systematically biased and is related to any other confounders such as age or sex. Understanding how reliable a set of cellular heterogeneity variables are for any individual sample is of increasing importance, as the interest in quantifying cellular composition has moved beyond just adjusting for it in epigenome-wide association studies, with these estimates also being analysed as variables of interest in their own right (Hannon et al., 2021a, Koestler et al., 2017, Wiencke et al., 2017).

In this study, we propose an accuracy metric that quantifies the **CE**ll **TY**pe deconvolution **GO**odness (CETYGO) score of a set of cellular heterogeneity variables derived from a genome-wide DNA methylation profile for an individual sample. While our method is applicable to any reference based deconvolution algorithm, and any reference panel of cell types, to demonstrate the utility of our approach we limit our characterisation to the Houseman algorithm and panels of blood cell types, which represent the majority of applications. We demonstrate that CETYGO indexes the accuracy of the prediction of cellular composition with simulations in which we manipulated the performance of the deconvolution. We then profile the statistical properties of CETYGO by applying it to a number of empirical datasets, to provide guidance on how it can be incorporated into whole blood DNA methylation studies. Finally, we use the CETYGO score to determine if they are any biases in the effectiveness of existing blood cell type reference panels. To enable the wider research community to incorporate our proposed error metric into their analyses, we have provided our methodology in an R package, CETYGO, as well as adding functions to the wateRmelon package.

## Materials and Methods

### Mathematical derivation of the CETYGO score

The DNA methylation profile of a bulk tissue can be defined as the sum of DNA methylation levels measured in the constituent cell types weighted by the proportion of total cells represented by that cell type. Mathematically we can represent this as

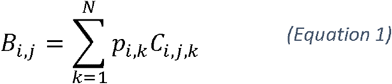

where

- *B_ij_* represents the DNA methylation level in the bulk tissue for sample i at site j
- *p_ik_* represents the proportion of cell type k in sample i
- *C_ijk_* represents the DNA methylation level for sample i at site j in cell type k, for N different cell types.

Typically in an epidemiological study, only the bulk tissue DNAm profile (*B_i,j_*) is measured. However, as cellular composition is an important confounder, it is desirable to know or estimate *p_i,k_* for all (major) cell types. Methods for this purpose, such as Houseman’s constraint projection approach, have been proposed that take advantage of reference profiles (i.e. *C_ijk_*) available to the research community to enable them solve for the unknown *p_i,k_.* This is achieved by selecting *M* DNA methylation sites that are highly discriminative of the cell types we want to estimate the proportions of. By definition, these sites exhibit low variation across individuals, and therefore it does not theoretically matter that we have not measured them in the same samples that we have bulk profiles from. If the estimated cell proportions (denoted 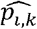) are accurate then the expected bulk tissue profile given this composition of cell types should closely resemble the observed data. We can substitute our estimated cell proportions, 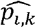, back into Equation 1, to calculate the expected profile of DNA methylation values (Equation 2).

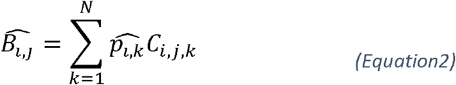

We define our error metric, CETYGO, as the root mean square error (RMSE) between the observed bulk DNA methylation profile and the expected profile across the *M* cell type specific DNA methylation sites used to perform the deconvolution, calculated from the estimated proportions for the *N* cell types (Equation 3). By definition, 0 is the lowest value the CETYGO score can take and would indicate a perfect estimate. Higher values of the CETGYO score are indicative of larger errors and therefore a less accurate estimation of cellular composition.

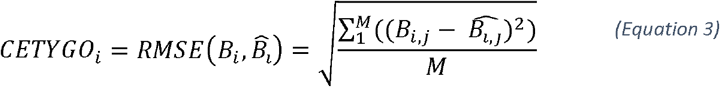

### Purified blood cell type reference panels

Genome-wide DNA methylation profiles for purified blood cell types generated using the Illumina 450K and EPIC microarray were obtained via the *FlowSorted.Blood.450k* and *FlowSorted.Blood.EPIC* R packages and formatted into matrices of beta values using commands from the *minfi(Aryee* et al., 2014) R package. From the 450K reference panel, we selected the six blood cell types that are mostly commonly used (B-cells, CD4+ T-cells, CD8+ T-cells, granulocytes, monocytes and natural killer cells) which were purified from whole blood from 6 male individuals using flow cytometry (Reinius et al., 2012). The EPIC reference panel contains profiles from antibody bead sorted neutrophils (n = 6), B-cells (n = 6), monocytes (n = 6), natural killer cells (n = 6), CD4+ T-cells (n = 7), and CD8+ T-cells (n = 6) (Salas et al., 2018). Prior to training any deconvolution models, both reference datasets were filtered to only include autosomal DNA methylation sites.

### Generation of deconvolution models and simulated whole blood profiles

To test the performance of CETYGO against a known truth, we trained a series of Houseman constraint projection deconvolution models and tested these against reconstructed whole blood DNA methylation profiles where we combined cell-specific profiles in a weighted linear sum of pre-specified proportions of each cell type. Depending on the specific testing framework, the training data comprised of all available samples that were not selected to be part of the testing data, such that the train and test data consisted of distinct sets of samples. It should be noted though, that in some scenarios they were from the sample experimental batch, and plausibly share technical batch-specific effects. We modified the *minfi* approach for implementing Houseman’s constrained projection methodology to omit the step within *estimateCellCounts(*) where the training and test data are normalised together, in order to explore the effect of normalization. This adaptation means that the cellular deconvolution and CETYGO calculation can be applied directly to a matrix of beta values, rather than requiring the raw data stored in an RGSet object. This makes it straightforward and computationally efficient to apply new reference panel (or include a new error metric) to an existing dataset. Briefly, our implementation performs an ANOVA to identify sites that are significantly different (p value < 1×10^-8^) between the blood cell types, selecting 100 sites per cell type (50 hypermethylated and 50 hypomethylated). These sites are then used to solve Equation 1 using quadratic programming, in essence a least squares minimisation, with the constraint that the proportions are greater than or equal to 0 and the sum of the proportions is less than or equal to 1.

In the first simulation analysis, we had six different combinations of training and testing data; within each reference panel (450K and EPIC), across reference panels without normalisation (450K to EPIC and EPIC to 450K) and across reference panels after stratified quantile normalisation as implemented in *minfi* of the combined training and test dataset (450K to EPIC and EPIC to 450K). To construct whole blood profiles for testing we isolated one sample of each cell type. When testing samples were selected from the 450K reference data, we selected a single individual as the test case and took all their purified samples, and therefore there were a maximum of 6 testing iterations (as there are 6 individuals). When testing samples were selected from the EPIC reference data, we randomly selected a test sample for each cell type (as they do not come from the same set of individuals), and repeated this process 10 times to get multiple sets of test data. We constructed whole blood profiles as a linear sum of these cell-specific profiles in a fixed ratio and a defined proportion of noise. Specifically,

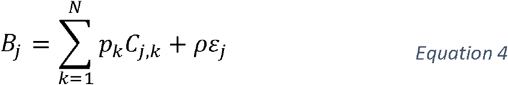

Where

- *B_j_* represents the simulated DNA methylation level in the bulk tissue at site j.
- *p_k_* represents the proportion of cell type k which were standardized for these series of simulations to the mean proportions reported in Reinius et al. (Reinius et al., 2012) (**Supplementary Table 1**).
- *C_j,k_* represents the DNA methylation level from the test sample for in cell type k at site j.
- ρ is the proportion of ‘noise’ and took the values 0,0.01,0.02,…,1,0.12,0.14,…0.5.
- ε_j_ is a random variable taken from a uniform distribution bounded by 0 and 1.
- 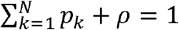

In total 31 simulated ‘noisy’ blood profiles were tested for each iteration of deconvolution model.

In the second simulation analysis, we focused on a single reference panel, the 450K reference panel. Here we tested a series of deconvolution models, where each cell type was omitted in turn from the reference panel, prior to training the model. Each of these leave one out models, was then tested against simulated whole blood profiles constructed from all six cell types. The five cell types included in the training data were again combined in fixed ratios calculated from the mean proportions reported by Reinius et al (**Supplementary Table 1**), with the omitted cell type included at increasing proportions (0.1,0.2,.,0.9). We used the same process to select testing samples as described before meaning that each of the leave one out models was tested against 9 simulated whole blood profiles in 6 different train test permutations.

In the third simulation analysis, we again focused on a single reference panel, the 450K reference panel. Here we tested all possible deconvolution models, containing between 3 and 5 of the 6 blood cell types, a total of 41 combinations. This time we tested the full spectrum of whole blood profiles in 0.1 units, where each cell type represented at least 0.1. In total 126 possible profiles were generated.

### Profiling the performance of CETYGO in real datasets

A summary of the 17 datasets used to profile CETYGO is provided in **Supplementary Table 2**. Datasets 2-9, 14, and 15 were generated by our group at the University of Exeter (www.epigenomicslab.com) have been previously published. The pre-processing and normalisation of these datasets is as described in the corresponding manuscripts. Datasets 1 and 16 were also generated by our group and are currently unpublished. They followed a standard QC pipeline and were normalised using *dasen*() in the *wateRmelon* package (Pidsley et al., 2013). Datasets 10-13 and 17 are publically available datasets obtained from GEO (https://www.ncbi.nlm.nih.gov/geo/). These data were put through a quality control pipeline which included checking the quality of the DNA methylation data (signal intensity, bisulfite conversion and detection p-values) prior to normalisation using *dasen*() in the *wateRmelon* package (Pidsley et al., 2013). For all datasets cellular deconvolution and the calculation of CETYGO was applied using a model trained with all samples for 6 cell types from the 450K reference panel.

To characterise the relationship between data quality metrics and CETYGO, we used an expanded version of Dataset 3 which retained the samples that failed quality control for either a technical or biological reason (n = 725). For this data we imported the raw signal intensities from the idat files for all samples using the *wateRmelon* package (Pidsley et al., 2013). Signal intensities for each sample were summarised as the median methylated (M) and unmethylated (U) intensity across all sites. Bisulfite conversion efficiency was calculated as the median beta value across 10 fully methylated control probes and converted to a percentage. Samples were then processed through *pfilter*() using the default settings. A sample was classed as a technical failure if either median signal intensity metric was less than 500, the bisulfite conversion statistic was less than 80% or it failed *pfilter().* In total 62 samples were classed as technical failures. Note these thresholds may not match up with the thresholds implemented in the quality control pipeline described in the original manuscript. All 725 samples were then normalised using *dasen* and cellular deconvolution and their CETYGO score estimated.

In order to test the effect of normalising the reference panel DNA methylation dataset (i.e. training data) with the bulk tissue dataset (i.e. the test data) we imported the raw signal intensities for Dataset 1. We the re-normalised these data in conjunction with the reference panel prior to performing cellular deconvolution and the calculation of CETYGO. To facilitate this we have adapted the *estimateCellCounts*() function in *minfi* (Aryee et al., 2014) to a new function *estimateCellCountsWithError*() which calculates CETYGO alongside performing the reference-based deconvolution. These values of CETYGO were compared to CETYGO calculated as described above using the dasen normalised betas, that were not normalised with the reference panel.

### Ethical approval

The study was approved by the University of Exeter Medical School Research Ethics Committee (reference number 13/02/009).

### Data and code availability

The DNAm data used in this study are available as R packages or via GEO (see **Supplementary Table 2** for details). We have provided the code for calculating the CETYGO score as an R package available via GitHub (https://github.com/ds420/CETYGO). The code to reproduce the analyses in this manuscript using our R package are also available via GitHub (https://github.com/ejh243/CETYGOAnalyses).

## Results

### CETYGO indexes the accuracy of cellular composition estimates in whole blood

The objective of this study was to define, validate and characterise a novel metric that can be used to assess the accuracy of DNAm-based cellular deconvolution in an individual sample. The CETYGO score captures the deviation between the observed DNAm profile and the expected profile for the given set of estimated cell type proportions, where values close to 0 indicate accurate estimates of cellular composition.

In order to test whether our proposed error metric CETYGO successfully captures inaccurate cellular heterogeneity estimates, we manufactured a series of bulk whole blood profiles where the cellular composition was known and could be estimated with varying degrees of accuracy. This was achieved by standardizing the ratios of the constituent blood cell types and adding an increasing proportion of random ‘noise’, which could reflect either biological variation, technical artefacts or imprecision in the assay (see **Materials and Methods**). The hypothesis is that as the proportion of noise increases, the estimation of cellular composition will be less accurate and the CETYGO score should correlate with the proportion of noise in the whole blood sample. To confirm that our simulation framework was fit for purpose, we calculated the RMSE between the fixed cell type proportions used to construct the whole blood profiles and the predicted values, observing that profiles with a higher proportion of noise were characterized by larger deviations from the truth (**Figure 1A**). Having manufactured a spectrum of inaccurate deconvolutions, we were able to determine whether the CETYGO score changed as a function of noise, finding that it successfully indexed accuracy with a monotonic relationship between the proportion of noise in a bulk sample and the CETYGO score (**Figure 1B**). We observed that for small proportions of noise (between 0 and 0.05) the accuracy estimates don’t vary very much, but once the proportion of noise goes above 0.05, the effect of additional noise on accuracy starts to accumulate. We also found that when the predictions were less accurate, the total sum of all estimated cell types for a sample was less than one and decreased as noise increased (**Figure 1C**).

**Figure 1.**
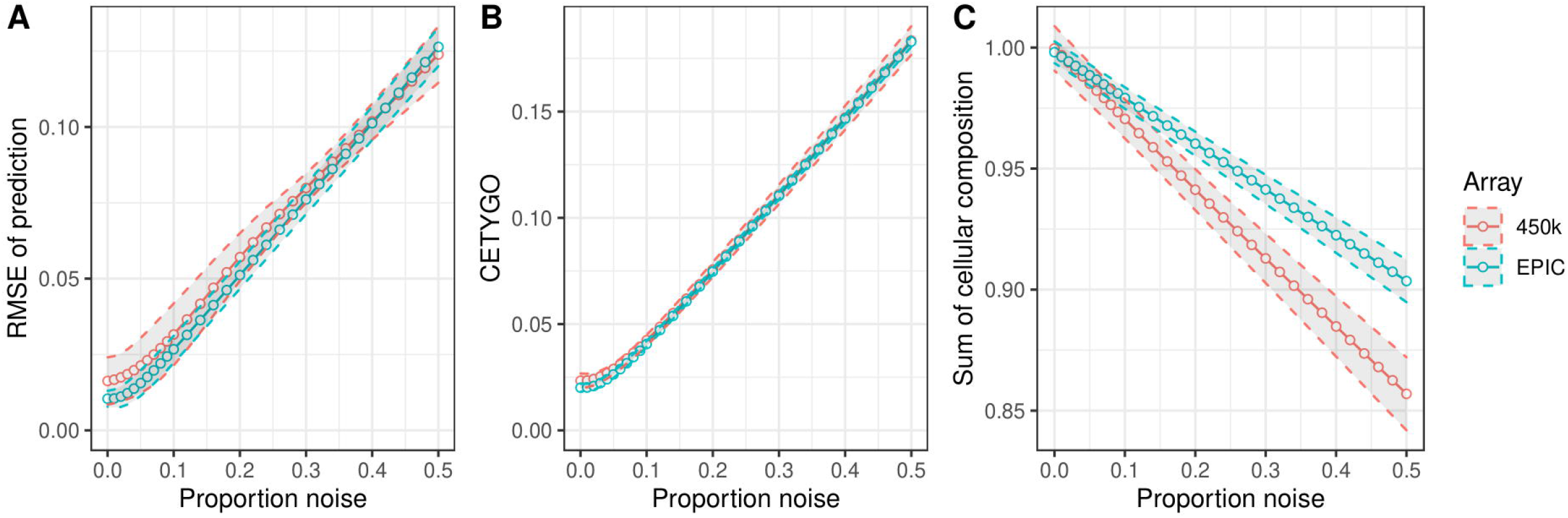
CETYGO captures variation in accuracy of cellular deconvolution in whole blood. Line graphs plotting the error associated with estimating the cellular proportions of reconstructed whole blood profiles with increasing proportion of noise (x-axis). Where the y-axis presents **A**) the root mean square error (RMSE) between the fixed cellular proportions used to construct the whole blood profiles and the estimated proportions generated with Houseman’s method, **B**) the error metric CETYGO and **C**) the sum of all proportions estimated. The points represent the mean value and the dashed lines the 95% confidence intervals calculated across multiple simulations. The two lines represent simulations constructed from reference data generated from two different platforms, the Illumina 450K and EPIC BeadChip microarrays.

In our simulation framework, we tested two independent reference datasets (Reinius et al., 2012, Salas et al., 2018), generated using different versions of the Illumina BeadChip array and incorporating subtly different panels of cell types (either granuloctyes or neutrophils). We subsequently repeated the simulation framework, but this time training the model using one reference panel (either 450K or EPIC) and testing it in simulations formulated from the other reference panel. This would allow us to explore how batch and normalisation strategy influences the accuracy of cellular deconvolution. These results showed the same general pattern across the different train-test pairings, where the CETYGO score captured decreasing accuracy in estimates of cellular composition (**Supplementary Figure 1**). Differences between datasets did lead to slightly increased imprecision at lower proportions of noise, but this scenario is arguably more representative of the typical application of cellular deconvolution algorithms, where the reference panel and bulk tissue test data are generated in different laboratories. Interestingly, we observed that when the training data was generated with the 450K array and applied to simulated bulk data generated from the EPIC array, the deconvolution was marginally more accurate potentially indicative of reduced signal-to-noise with the EPIC array. In general, whether the two batches of data were normalised together or not had a minimal effect on deconvolution accuracy, measured by either RMSE (**Supplementary Figure 1A**), or the CETYGO score (**Supplementary Figure 1B**), There was however, subtle variation dependent on which panel was used as the training data, suggesting that technology, data quality or cell purity is more important than normalisation strategy. Given the comparable performance of the two reference panels, all subsequent analyses were performed with the 450K reference panel only.

### CETYGO is inflated when applied to incomplete cellular reference panels

Another scenario where inaccurate deconvolutions are likely to occur is when the reference panel of cell types for deconvolution is incomplete. One of the constraints set when implementing Houseman’s method to solve for cellular composition proportions is that the sum of the proportions of the cell types in the panel ≤ 1. In other words, all the cells present in the bulk tissue are (virtually) completely represented by the cell types in the reference panel. When an abundant cell type is missing due to lack of reference data, theoretically, this may lead to errors, as the unrepresented proportion of the bulk tissue will need to be (incorrectly) assigned to an alternative cell type. To explore this, we dropped each cell type in turn from the reference panel, and recalculated the cellular proportion estimates for reconstructed whole blood profiles that included the missing cell type, in increasing proportions. We found that the CETYGO score had a monotonically increasing relationship with the true proportion of the missing cell type (**Figure 2**). Of note, the magnitude of the CETYGO score in blood data depended on which blood cell type was missing, with the omission of B-cells, leading to the largest errors and the omission of CD8+ T-cells the smallest effect. This is likely due to the methylomic similarity of the two sets of T-cells, whereby CD4+ T-cells are a good alternative to CD8+ T-cells, and suggests that at sites included on the 450K array, B-cells have the most distinct profile. We expanded this framework further to omit up to 3 cell types from the training model, finding that the CETYGO score generally decreases as both the number of cell types in the model increases and the proportion of cells represented in the model increases (**Figure 3**). However, the distributions of the CETYGO score across different panels of cell types applied to different compositions of whole blood are overlapping and have long tails, highlighting that there are some scenarios where a model with 3 cell types, outperforms a model with 4 or 5 cell types dependent on the abundance of each cell type in the bulk tissue.

**Figure 2.**
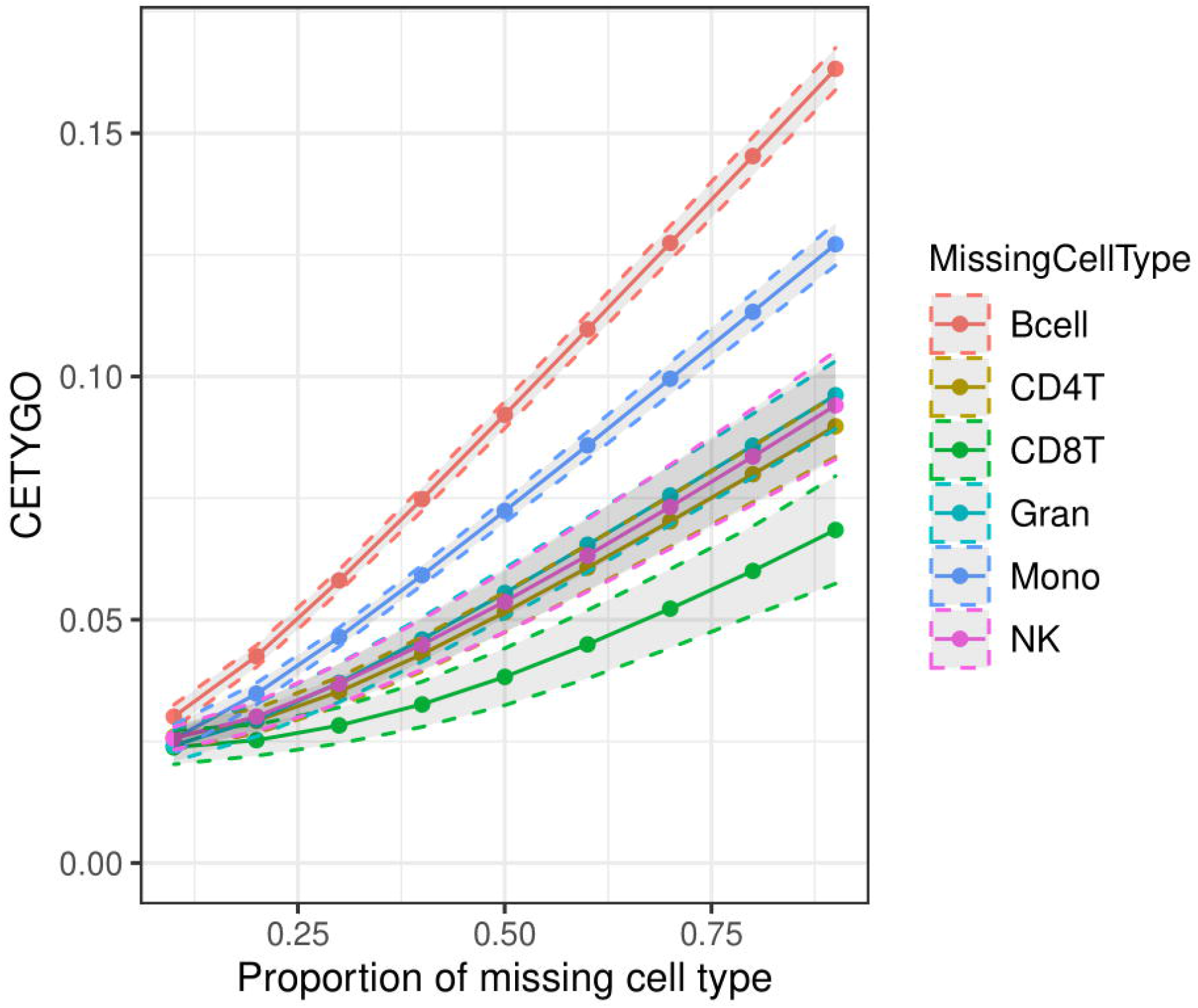
Cell type dependent effects on accuracy when omitted from reference based cellular deconvolution algorithms. Line graph of the error associated with estimating the cellular proportions of reconstructed whole blood profiles where the reference panel is missing one of six cell types. Each coloured line represents a different cell type being omitted from the reference panel, but included in the reconstructed whole blood profiles used for testing. Plotted is the proportion in the testing profile that the missing cell type is set to occupy (x-axis) against the error, measured using CETYGO, of the deconvolution (y-axis). The points represent the mean value and the dashed lines the 95% confidence intervals calculated across multiple simulations.

**Figure 3.**
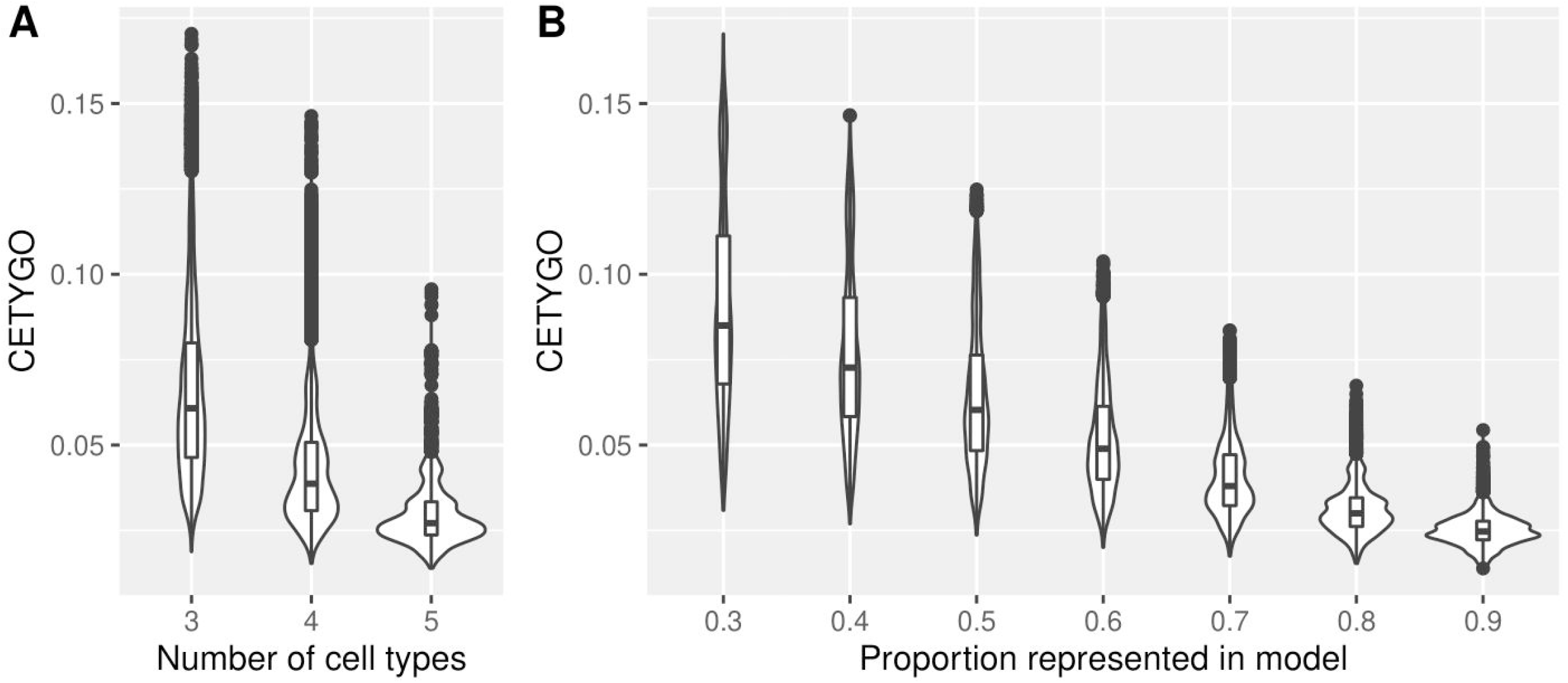
The accuracy of cellular heterogeneity estimation increases as the reference panel becomes more representative. Violin plots of the error associated with estimating the cellular proportions of reconstructed whole blood profiles where the reference panel is missing between one and three cell types. Each violin plot shows the distribution of the error, measured using CETYGO, of the deconvolution (y-axis) grouped by **A**) the number of cell types included in the reference panel and **B**) the proportion of cells in the reconstructed whole blood profile that are from cell types included in the reference panel.

### CETYGO distinguishes nonsense applications

Having demonstrated the sensitivity of the CETYGO score to detect noisy and incomplete estimates of cellular heterogeneity, we next tested its behaviour when applied to real data in order to provide guidance to the wider research community about how it can be interpreted in the context of epidemiological studies. To this end, we estimated the cellular proportion of six blood cell types and the CETYGO score associated with the estimation for 10,447 DNA methylation profiles, across 17 different datasets and 17 different sample types (**Supplementary Table 2**). 7,184 (68.8%) of these represent realistic applications as the profiles were derived from blood tissue types and can be used to infer the expected distribution of CETGYO scores across a range of experimental and biological sources. The remaining 3,263 (31.2%) represented “nonsense” applications as these profiles were generated from non-blood samples and can be used to highlight whether the CETYGO score can distinguish sensible deconvolutions. In general, there was a clear dichotomy between the output for these two types of sample; CETYGO scores for blood samples were typically < 0.1 and CETYGO scores for non-blood tissues were > 0.1 (**Figure 4**). The median CETYGO score across all whole blood samples was 0.0524 (inter-quartile range = 0.04550-0581). Within the whole blood samples there was a bimodal distribution, which on closer inspection was driven by platform, with datasets generated with the 450K array associated with lower CETYGO scores than those generated using the EPIC array (**Supplementary Figure 2**). Limiting our comparison to Dataset 8 where we had matched whole blood and purified blood cell types from the same individuals (Hannon et al., 2021b), we observed that purified blood cell types were predicted with higher error than whole blood (**Supplementary Figure 3**), with significant differences for all cell types, bar granulocytes (**Supplementary Table 3**). This suggests that it is more challenging to determine a cell type is pure, than to deconvolute a mixture of cell types. We also noted that the CETYGO score was significantly higher for both cord blood (mean difference = 0.0207; T-test p-value < 3.42×10^-363^) and neonatal blood spots (mean difference = 0.0307; T-test p-value = 9.19×10^-62^) compared to whole blood. This is in agreement with previous studies suggesting that the standard panel of major blood cell types is not the most appropriate for the assessment of cellular heterogeneity in blood samples obtained for neonatal epigenetic studies (Bakulski et al., 2016).

**Figure 4.**
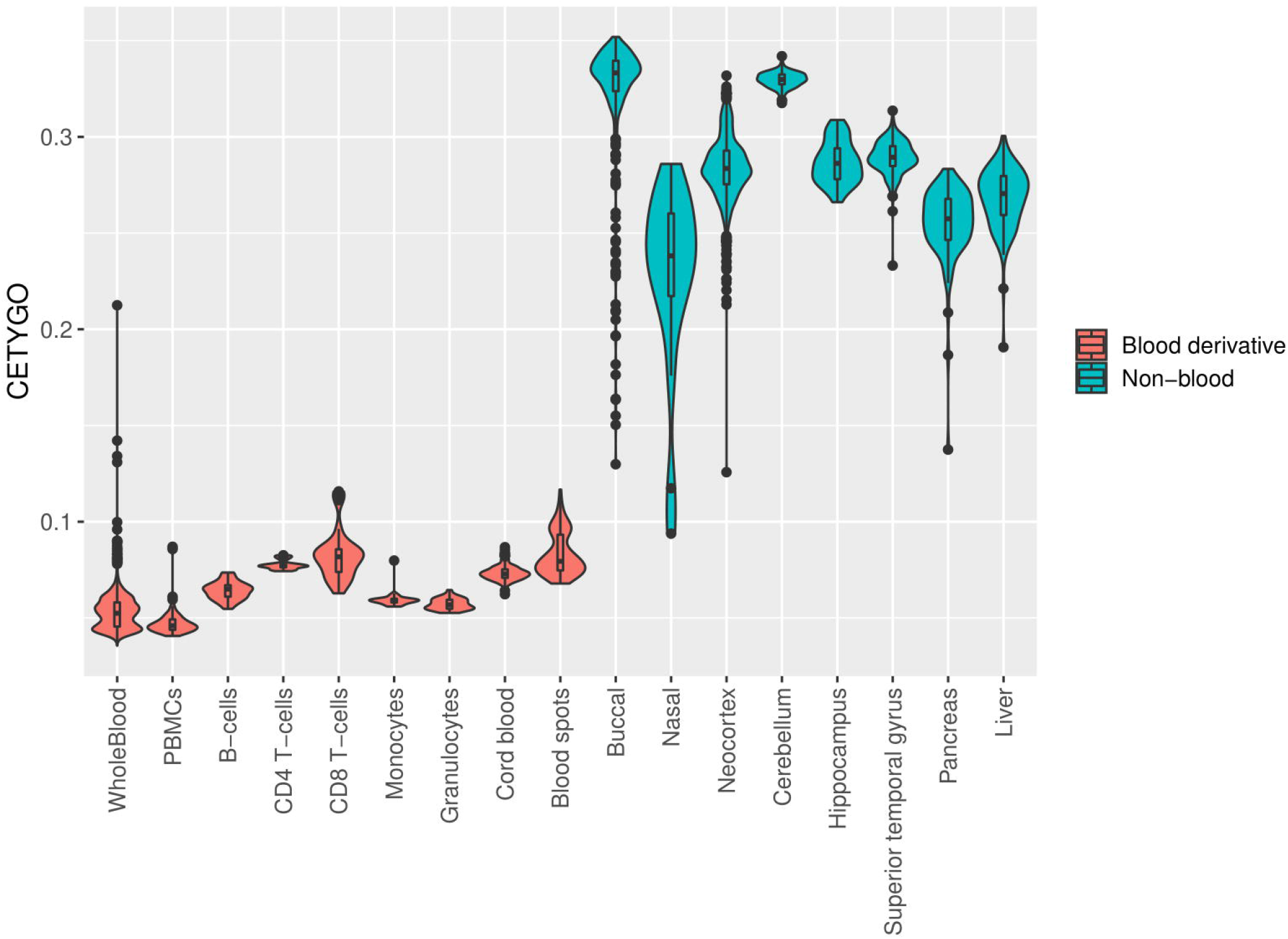
CETYGO captures the tissue specificity of deconvolution reference panels. Violin plots of the error associated with estimating the cellular proportions where a reference panel consisting of six blood cell types was applied to 10,447 DNA methylation profiles, across 18 different datasets and 20 different sample types. Each violin plot shows the distribution of the error, measured using CETYGO, of the deconvolution (y-axis) grouped by the tissue/cell-type, where the violins are coloured to highlight which samples are derived from blood, which are human derived non-blood bulk tissue, and which are human derived cell-lines.

### Cellular heterogeneity estimates are biased by technical factors

While the distribution of CETYGO score across whole blood samples was fairly narrow, we wanted to explore whether CETYGO scores could be used to detect biases in the estimation of cellular composition from whole blood DNA methylation profiles. In the simulation study we showed that noisy DNA methylation profiles lead to less accurate estimates of cellular composition. In real data, technically noisy signals should be excluded as part of the pre-processing pipeline in order to improve the power to detect differences between groups. We hypothesized that samples excluded based on technical quality metrics are likely to have higher deconvolution errors as measured by the CETYGO score. Comparing CETYGO scores against standard quality control metrics we found that higher values of the CETYGO score were associated with lower median signal intensities and lower bisulfite conversion statistics (**Supplementary Figure 4**), consistent with our hypothesis.

The vast majority of DNA methylation studies perform normalisation to align the distributions across samples, and ultimately make the data more comparable, particularly where data have been generated across multiple batches. We hypothesised that normalising reference data and test data together to make the genome-wide profiles more similar would attenuate the discriminative signals between cell types and negatively affect the performance of cellular deconvolution. We therefore compared the CETYGO scores calculated with and without normalisation of the test data with the reference panel for Dataset 1. In general, the overall distribution of values did not differ dramatically between normalisation strategies.

However, we did observe that when the reference panel (which is all male) was normalised with the test data, there was a clear bias towards females having higher error (**Supplementary Figure 5**), consistent with analyses showing that normalisation can introduce sex effects(Wang et al., 2021). In contrast, our adapted method, where we normalised the data separately, was characterized by a dramatically reduced sex difference.

### Cellular heterogeneity estimates are biased by age, sex and smoking status

Across the 6,351 whole blood samples included in our analysis we fitted a linear regression model to test the influence of additional factors on CETYGO scores (**Supplementary Table 4**).As well as the platform effects we described earlier (p-value = 2.72×10^-223^) there were further significant differences between datasets (p-value = 1.75×10^-222^) even after controlling for platform. We also found that every biological factor we tested had a significant association with CETYGO (**Supplementary Figure 6**). This included a negative association with age (coefficient = −7.1×10^-5^, p-value = 0.00215), a positive association with age squared (coefficient = 8.8×10^-7^, p-value = 0.000189), sex (mean difference in males = 9.6×10^-4^, p-value = 4.03×10^-12^) and a positive association with smoking score (coefficient = 6.7×10^-5^, p-value = 1.84×10^-6^).

### Inaccuracies in DNA methylation prediction algorithms are concordant across predictors for different phenotypes

Finally, we were interested in whether inaccuracy in cellular deconvolution was mirrored by inaccuracies in other epigenetic predictors. Comparing CETYGO against the deviation between chronological age and epigenetic age predicted with the Horvath multi-tissue clock (Horvath, 2013), we found a significant positive relationship (coefficient = 43.0, p-value = 1.68×10^-5^) highlighting that samples with inaccurate cellular deconvolution have a larger difference between epigenetic age and chronological age (**Figure 5**).This suggests that studies which use the residual between epigenetic age and chronological age as a proxy for accelerated aging are potentially just modelling the imprecision in the technology.

**Figure 5.**
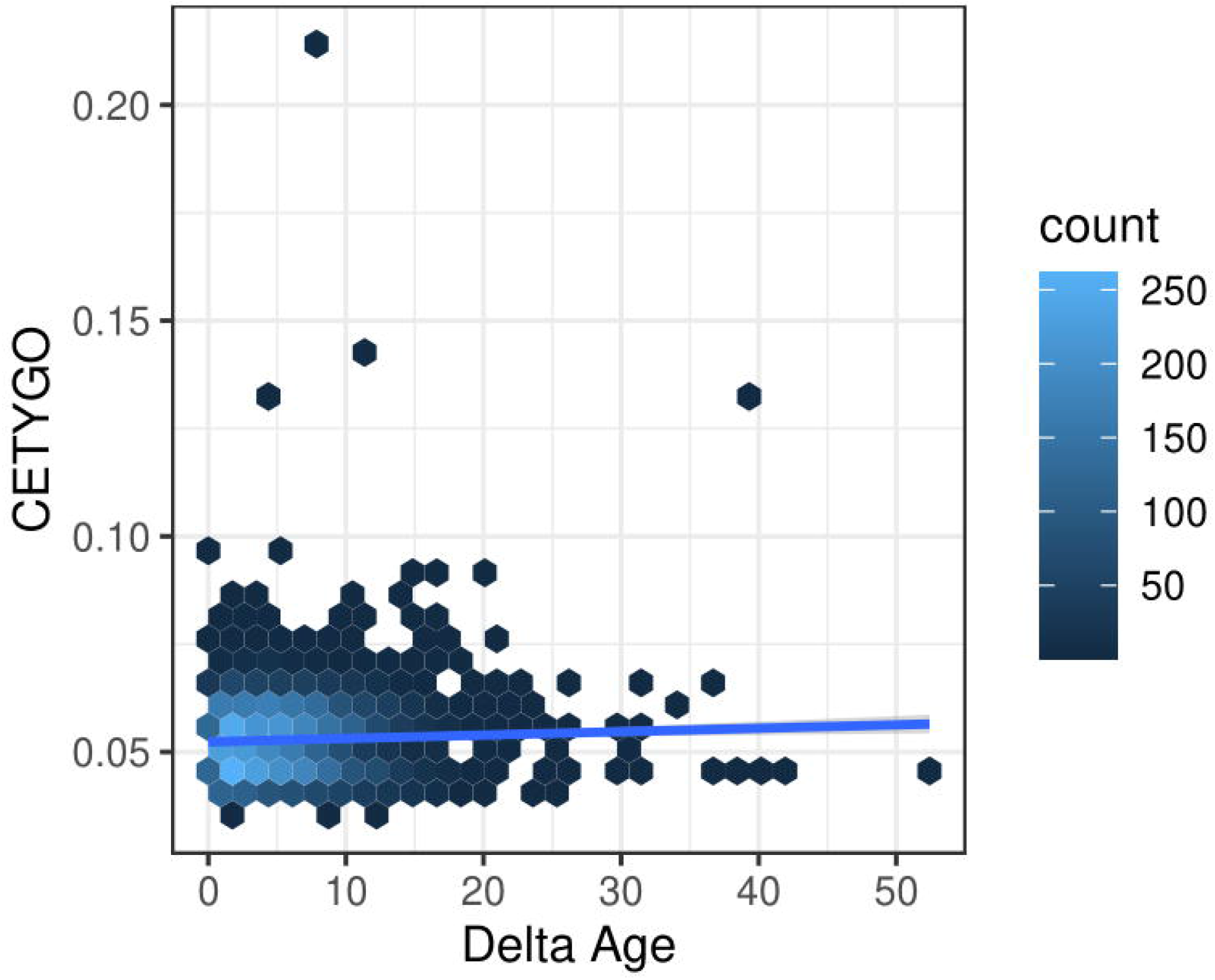
Error in estimation of cellular heterogeneity from DNA methylation data correlates with error from epigenetic clock algorithms. Heatscatterplot of the error measured using CETYGO (y-axis), associated with estimating the cellular proportions across 6,351 whole blood profiles against the difference between the sample’s chronological age and age predicted using Horvaths pan-tissue algorithm from the DNA methylation data (Delta age; x-axis). The colour of the points represents the density of points at that location.

## Discussion

The estimation of cellular composition is vital in epigenetic epidemiology, with these variables being included as co-variates in analyses to minimise the effect of confounding. To compliment these analyses, we have described and validated a novel error metric – CETYGO - that enables the *accuracy* of the deconvolution to be quantified at an individual sample level. Our results demonstrate that the CETYGO score consistently distinguishes inaccurate and incomplete deconvolutions when applied to reconstructed whole blood profiles and support its inclusion in future DNA methylation association studies to identify scenarios, or individual cases, when cell composition estimates are unreliable. We have applied it to several existing datasets to further characterise the performance of the predominant application with a reference panel of blood cell types. These analyses provided a number of insights. First, our results indicate that cell types are not equal when it comes to deconvolution accuracy. For example, the omission of B-cells from the standard blood reference panel had the most dramatic effect on their accuracy, while the omission of one of the two types of T-cells had the smallest effect. This is consistent with previous reports that the DNA methylation profile of B cells is relatively distinct to that of other blood cell-types, with the profiles of the two T-cells being most similar (Hannon et al., 2021b). Second, we highlighted that the estimation of cellular deconvolution using the existing reference panel is biased. Specifically, it is less accurate in females, neonates, older individuals and smokers. This has important consequences for epigenome-wide association studies, as it may indicate that existing efforts to adjust for cellular heterogeneity may be less effective in some sets of samples. This emphasizes the need to thoroughly benchmark all reference panels and characterise which scenarios they are appropriate for and to increase the diversity of available reference panels.

Our primary motivation was to develop a metric that that could be used to assess for an individual sample, how reliable derived estimates of cellular heterogeneity are. To facilitate this we have calculated the CETYGO score in >6,300 whole blood profiles, and provided some guidance about how to interpret the metric. Our data suggest that a CETYGO score > 0.1 is consistent with the reference panel not being relevant for the specific tissue being profiled. Although incorrect tissue, had the most dramatic effect, we also found that elevated CETYGO can be induced by poor quality DNAm data, where the noise to signal ratio is elevated, generating less sensitive DNA methylation profiles to the extent that it interferes with the accuracy of the deconvolution model. This can be mitigated by implementing stringent pre-processing pipelines to remove poor quality data. In particular, the principle behind our metric is comparable to the quality control metric DMRSE available in the wateRmelon R package(Pidsley et al., 2013). However, even within the pre-processed datasets used in our study there were a handful of samples with outlier CETYGO values. For this reason, we suggest that CETYGO should be added to existing pipelines to provide confidence in analyses that incorporate cellular composition variables. To facilitate this, we have made our method available as a standard alone R package – CETYGO - available via GitHub which adapts the existing workflow within minfi (Aryee et al., 2014) to simultaneously calculate the CETYGO score alongside the estimation of cellular composition variables using Houseman’s algorithm. In this way it can easily be adapted for use with other available reference panels, both now and in the future. We have also integrated the CETYGO score into the wateRmelon function *EstimateCellCounts.wmln(*), used to predict cell type composition, providing users with their deconvolution accuracy estimate when they predict composition.

Our findings should be considered in the light of a number of limitations. First, for the purpose of validation, we limited our analyses to the most commonly used deconvolution algorithm, Houseman’s constrained projection approach (Houseman et al., 2012), and the most commonly used bulk tissue, whole blood, for which a previously validated reference panels (Accomando et al., 2014, Koestler et al., 2013) exist. Comparisons of the different methodologies for inferring cellular heterogeneity estimates from bulk tissue have concluded that no single method is superior across all test scenarios (Teschendorff et al., 2017). Theoretically, though, the concept behind the CETYGO score should be extendable to any reference based deconvolution algorithm or reference panel of cell types and therefore applicable to any tissue, organism, or DNA methylation profiling technique and could be used to compare the performance of difference algorithms within a single dataset where true cellular heterogeneity is unknown. Second, our method assumes that the cell-specific sites used to estimate cellular composition are not influenced by any exposure. If differences were induced at these sites, this would cause the error to be overestimated. This assumption is also made by most deconvolution algorithms, and it has been suggested that it is unlikely to be a major concern (Teschendorff and Zheng, 2017). Third, we limited the majority of analyses to a reference panel generated with the 450K array and therefore, the conclusions regarding the effect of the specific blood cell types on accuracy may be influenced by the subset of genomic loci included on that technology.

In summary, we have proposed a new metric, CETYGO, to evaluate the accuracy of reference based cellular deconvolution algorithms at an individual sample level. We believe, this tool will be asset in studies of DNA methylation and have demonstrated how it can be used to assess bias in reference panels, and to identify unreliable estimates of cellular composition.

## Acknowledgements

We are grateful to Alice Franklin and Sim Lin for testing out the CETYGO package.

## Funding

D.S.V is funded by a BBSRC CASE PhD studentship. E.H is supported by an Engineering and Physical Sciences Research Council Fellowship EP/V052527/1. E.H., J.M., E.L.D, and L.C.S. were supported by Medical Research Council grant MR/R005176/1. G.S. was supported by a PhD studentship from the Alzheimer’s Society. The generation of the DNA methylation data was primarily funded by Medical Research Council grant MR/K013807/1. Data analysis was undertaken using high-performance computing supported by a Medical Research Council (MRC) Clinical Infrastructure award (M008924). For the purpose of open access, the author has applied a ‘Creative Commons Attribution (CC BY) licence to any Author Accepted Manuscript version arising.

## Disclosure of interest

The authors report no conflict of interest.

## Notes

https://www.ncbi.nlm.nih.gov/geo/query/acc.cgi?acc=GSE80417

https://www.ncbi.nlm.nih.gov/geo/query/acc.cgi?acc=GSE84727

https://www.ncbi.nlm.nih.gov/geo/query/acc.cgi?acc=GSE152027

https://www.ncbi.nlm.nih.gov/geo/query/acc.cgi?acc=GSE147221

https://www.ncbi.nlm.nih.gov/geo/query/acc.cgi?acc=GSE88890

https://www.ncbi.nlm.nih.gov/geo/query/acc.cgi?acc=GSE89705

https://www.ncbi.nlm.nih.gov/geo/query/acc.cgi?acc=GSE58885

https://www.ncbi.nlm.nih.gov/geo/query/acc.cgi?acc=GSE89702

https://www.ncbi.nlm.nih.gov/geo/query/acc.cgi?acc=GSE89703

https://www.ncbi.nlm.nih.gov/geo/query/acc.cgi?acc=GSE89707

### Competing Interest Statement

The authors have declared no competing interest.

