## Supplementary Figure for "Uncertainty quantification of reference based cellular deconvolution algorithms"

**Supplementary Figure 1. CETYGO captures variation in accuracy of cellular deconvolution in whole blood independent of experimental batch and normalisation strategy.** Line graphs plotting the error associated with estimating the cellular proportions of reconstructed whole blood profiles with increasing proportion of noise (x-axis). Where the y-axis presents **A**) the root mean square error (RMSE) between the fixed cellular proportions used to construct the whole blood profiles and the estimated proportions generated with Houseman’s method, **B**) the error metric CETYGO and **C**) the sum of all proportions estimated. The points represent the mean value and the dashed lines the 95% confidence intervals calculated across multiple simulations. The two lines represent simulations constructed from reference data generated from two different platforms, the Illumina 450K and EPIC BeadChip microarrays.

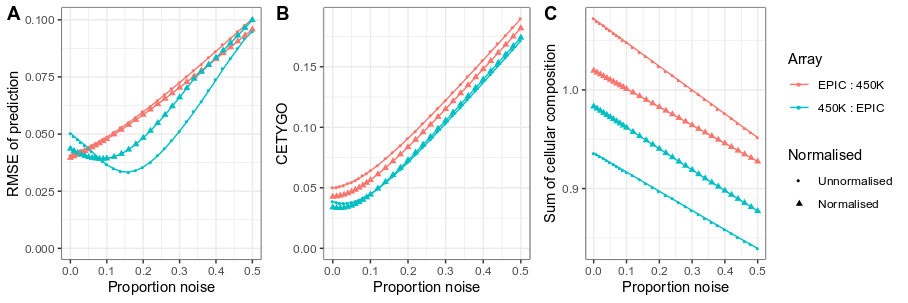

**Supplementary Figure 2. CETYGO varies across datasets.** Violin plots of the error associated with estimating the cellular proportions where a reference panel consisting of six blood cell types is applied to 10,447 samples taken from a wide range of DNA methylation datasets including a range of cell/tissues type. Each violin plot shows the distribution of the error, measured using CETYGO, of the deconvolution (y-axis) grouped by the dataset, where the violins are coloured to highlight which version of the Illumina Beadchip Methylation array was used.

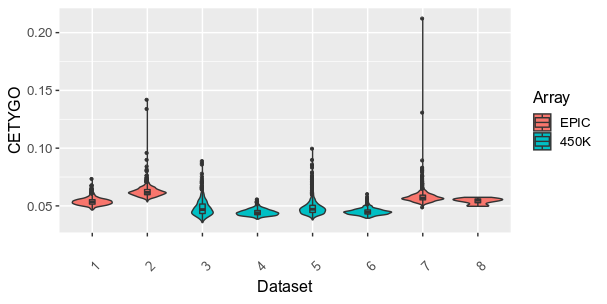

**Supplementary Figure 3. CETYGO is higher in purified cell types compared to bulk tissue.** Violin plots of the error associated with estimating the cellular proportions where a reference panel consisting of six blood cell types is applied to matched whole blood and purified blood cell types from the same individuals. Each violin plot shows the distribution of the error, measured using CETYGO, of the deconvolution (y-axis) grouped by the sample type.

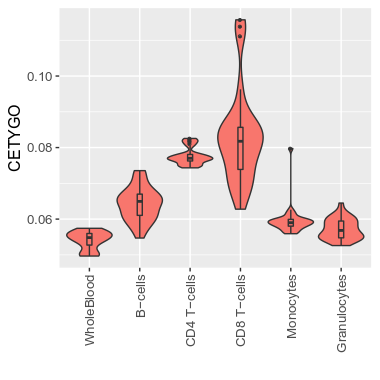

**Supplementary Figure 4. CETYGO correlates with metrics of data quality.** Summaries of the error associated with estimating the cellular proportions as a function of quantitative metrics of DNA methylation array signal for 725 samples from dataset 3. **A)** Violin plot of the distribution of CETYGO, grouped by whether the sample is of sufficient quality to pass the quality control pipeline. Scatterplots of the error, measured using CETYGO (y-axis) for each sample against, **B**) the median methylated (M) intensity across all sites on the microarray, **C**) the median unmethylated (U) intensity across all sites on the microarray, **D**) the bisulfite conversion % calculated as the mean across 10 fully methylated control probes. In panels, **B**, **C** and **D**, the points are coloured by whether the sample passed quality control in panel **A** or not.

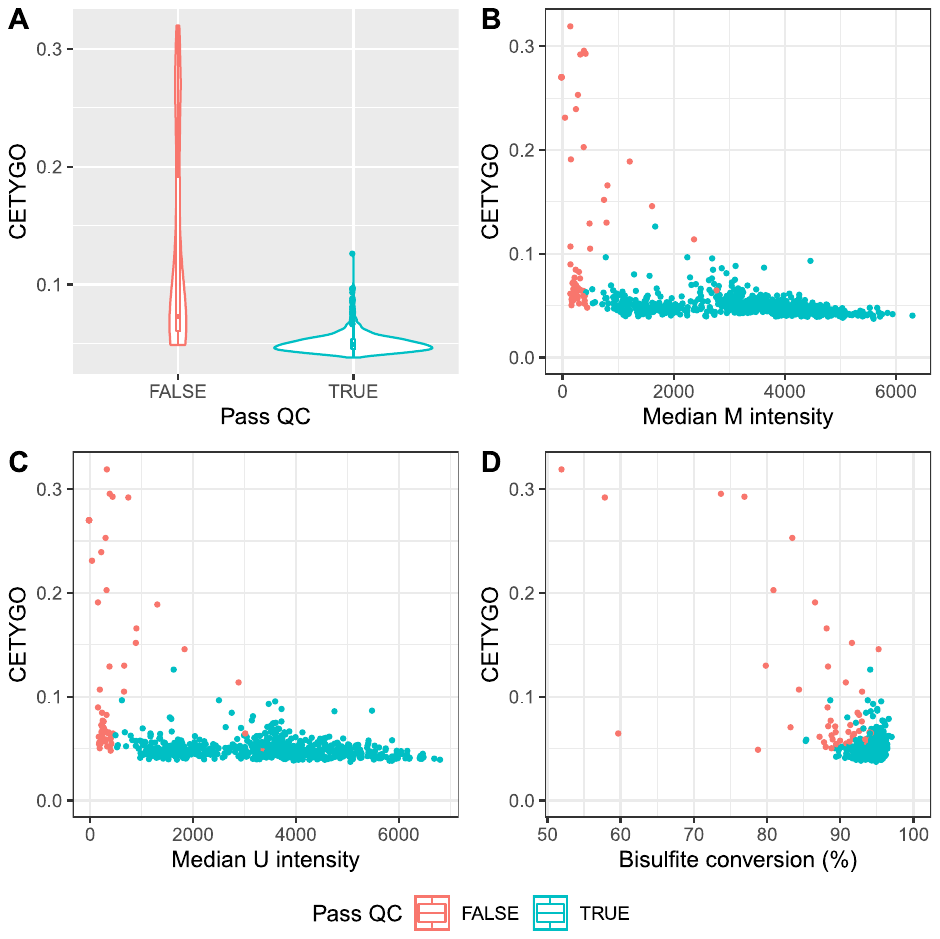

**Supplementary Figure 5. Normalisation of reference data and bulk test data induces sex effects on CETYGO.** Violin plots showing the distribution of the error associated with estimating the cellular proportions, measured by CETYGO (y-axis), grouped by sex where in panel the samples where the reference data and whole blood test samples were normalised **A**) together and **B**) separately for 1,234 samples from Dataset 1.

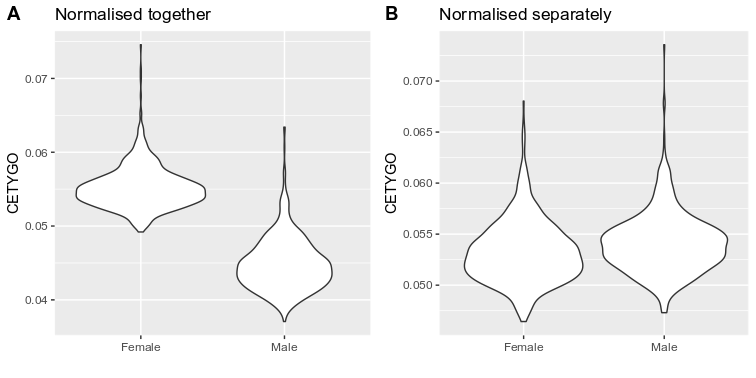

**Supplementary Figure 6. CETYGO indicates bias in existing blood cell type reference panels.** Summaries of the error associated with estimating the cellular proportions as a function of biology factors for 6,351 whole blood samples. **A**) Violin plots showing the distribution of the error associated with estimating the cellular proportions, measured by CETYGO (y-axis), grouped by sex and platform. Scatterplots of CETYGO (y-axis), against **B**) chronological age measured in years and **C**) DNA methylation derived smoking score. Points are coloured by the version of the Illumina BeadChip array used to generate the DNA methylation profile, and the lines are the regression lines generated with ggplot2 for each array platform.

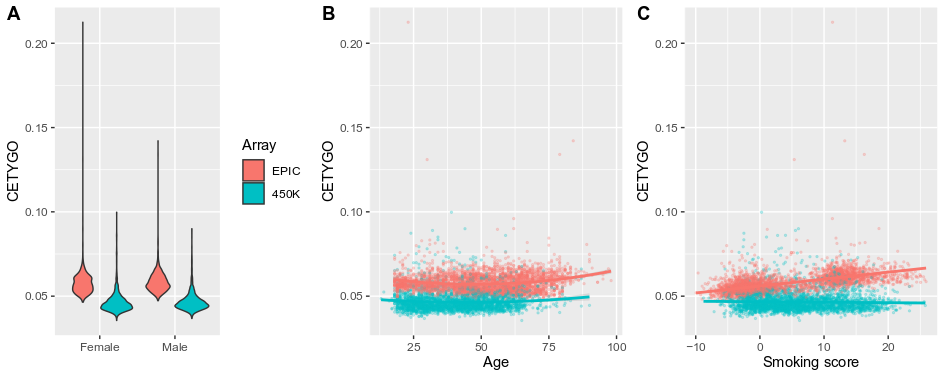
